# Differential PCDH1 usage by Laguna Negra and Black Creek Canal hantaviruses

**DOI:** 10.64898/2026.10.05.756634

**Authors:** Nahomi Guerra-Pilaquinga, Cierra Word, Upendra P. Lambe, Lohit Khera, Ridwan Arinola, Ramandeep Kaur, Maria Eugenia Dieterle, Rohit K. Jangra

**Author notes:** Corresponding author: Rohit K Jangra.

## Abstract

Laguna Negra virus and Black Creek Canal virus are hantaviruses associated with hantavirus cardiopulmonary syndrome (HCPS), but their cellular entry receptors have not been defined. We used replication-competent recombinant vesicular stomatitis viruses (rVSVs) bearing hantavirus Gn/Gc glycoprotein complexes to compare entry by these viruses, the Old World rodent-borne Dabieshan virus and Tigray virus, and the shrew-borne Cao Bang virus and Amga virus. All six Gn/Gc complexes supported infection of primary human pulmonary microvascular endothelial cells, with virus-specific efficiencies and kinetics. Genetic ablation of protocadherin-1 (PCDH1) reduced infection mediated by Laguna Negra virus and Black Creek Canal virus Gn/Gc in human osteosarcoma and endothelial cells, whereas PCDH1 re-expression restored susceptibility. Both viruses were captured by a soluble PCDH1 protein comprising the first two extracellular cadherin domains. Infection was also inhibited by a soluble decoy containing the first cadherin domain, and by an antibody specific to that domain, indicating that the first PCDH1 cadherin domain mediates viral engagement. Black Creek Canal Gn/Gc-bearing rVSVs incorporated less Gn/Gc and showed lower apparent capture by PCDH1 than Laguna Negra virus and Andes virus Gn/Gc-bearing particles, even after normalization for glycoprotein abundance. In contrast, PCDH1 depletion did not detectably alter infection mediated by the other four hantavirus Gn/Gc complexes. Thus, Laguna Negra virus and Black Creek Canal virus use PCDH1 for efficient entry but differ in apparent receptor engagement and responses to PCDH1-directed inhibitors, whereas all six viruses are potently neutralized by the broadly reactive Gn/Gc-directed antibody ADI-42898.

**Importance:** The receptors used by many medically important hantaviruses remain incompletely characterized. Using BSL2-compatible rVSVs bearing hantavirus glycoproteins, we show that Laguna Negra virus and Black Creek Canal virus, two hantaviruses associated with human cardiopulmonary disease, used protocadherin-1 (PCDH1) for efficient entry into human cells. PCDH1-directed inhibitors blocked infection, but the two viruses differed in apparent receptor engagement and inhibitor sensitivity, showing that PCDH1 usage is not uniform even among closely related viruses. Gn/Gc complexes from four additional divergent hantaviruses carried by shrews and Old World rodents infected primary human endothelial cells without detectable PCDH1 dependence, emphasizing the need to identify alternative entry factors. A broadly reactive antibody neutralized every virus in the study panel, supporting conserved viral glycoprotein surfaces as therapeutic targets when receptor usage is diverse or unknown.

## Introduction

Entry of enveloped viruses into host cells is a major determinant of tissue tropism and cross-species transmission(1). Mammalian hantaviruses (family *Hantaviridae*, order *Elliovirales*) are emerging zoonotic pathogens that cause severe, often fatal illness in humans after transmission from chronically infected small-mammal reservoirs, manifesting as hemorrhagic fever with renal syndrome (HFRS) in Eurasia and hantavirus cardiopulmonary syndrome (HCPS) in the Americas (2, 3). The public health significance of hantaviruses is underscored by their recurring emergence, including a multi-country HCPS outbreak in April-May 2026, which involved limited secondary transmission and resulted in three reported deaths (4–6). Despite significant morbidity and high case-fatality rates approaching 40% for HCPS, no FDA-approved prophylactics or antiviral therapeutics are currently available (7). Cellular entry of hantaviruses is orchestrated entirely by the envelope glycoprotein complex Gn/Gc (8, 9). Gn and Gc are synthesized as a single polyprotein precursor encoded by the medium (M) genomic segment, co-translationally cleaved by cellular signal peptidases, and assemble into spike-like tetramers of Gn/Gc heterodimers on the virion envelope (8–11)These glycoproteins coordinate entry: the membrane-distal Gn subunit likely mediates host receptor engagement, while the class II fusion protein Gc drives low-pH-dependent membrane fusion within late endosomes envelope (8–12) Despite the identification of dozens of hantavirus species across rodent and insectivore hosts globally, host entry receptors and neutralization vulnerabilities for most divergent species remain undefined (10, 13, 14).

Candidate hantavirus attachment factors and receptors initially identified through antibody blockade, ligand competition, or *in vitro* binding assays include *β1*, *β2*, and *β3* integrins, decay-accelerating factor (DAF/CD55), and gC1qR complement receptor (15–19). However, CRISPR/Cas9-mediated loss-of-function screens in human microvascular endothelial cells, a primary *in vivo* target cell type, demonstrated that knockout of *ITGB1, ITGB3,* or *DAF* had little or no effect on infection by a panel of virulent hantaviruses, indicating that these molecules are dispensable for endothelial cell entry (20). Unbiased haploid-genetic screening instead identified protocadherin-1 (PCDH1) as an essential entry receptor for Andes virus (ANDV, *Orthohantavirus andesense*) and Sin Nombre virus (SNV, *Orthohantavirus sinnombreense*): its membrane-distal extracellular cadherin domain 1 (EC1) directly binds Gn/Gc, and genetic ablation of *PCDH1* in cultured cells and Syrian hamsters protects against ANDV infection and lethal disease (21). Point mutations within the EC1 contact surface disrupt infection, establishing PCDH1-EC1 as the functional interface (22). Although PCDH1 usage has been established for multiple hantaviruses, the glycoprotein determinants that govern receptor recognition across divergent lineages remain incompletely defined (20–22). PCDH1 usage has also been documented beyond the canonical New World clade, including in vole-and mole-borne viruses (23), but its distribution across other rodent-and shrew-borne clades has yet to be fully mapped.

Gn/Gc amino acid sequences differ by roughly 40-60% between hantavirus lineages, and most glycoprotein-directed monoclonal antibodies (mAbs) raised or isolated against one species show little heterologous neutralization (24, 25). Consequently, broadly active entry inhibitors may need to target either conserved virus-receptor interfaces or rare, functionally constrained epitopes on the viral spike (11, 21, 24, 25). On the host side, the EC1-specific mAb-3305, soluble histidine-tagged PCDH1 ectodomain proteins containing EC1-2 or EC1-4 (sEC1-2, sEC1-4), and a human Fc-tagged soluble EC1 decoy (EC1-Fc) block PCDH1-Gn/Gc engagement and neutralize PCDH1-dependent viruses (21–23). Conversely, targeting conserved epitopes on the viral spike can provide broad neutralization across clades. In particular, mAb-42898, isolated from a Puumala virus (PUUV, *Orthohantavirus puumalaense*)-convalescent individual, binds a quaternary epitope bridging the Gn capping loop and Gc fusion loops, neutralizes multiple HCPS-and HFRS-causing viruses, and protects rodents against lethal challenge (24, 26). Evaluating an expanded panel of uncharacterized Gn/Gc complexes against both receptor-targeted and spike-targeted approaches can reveal conserved viral vulnerabilities and inform the development of broadly active therapeutics.

Functional entry studies of less-characterized hantaviruses are constrained by the limited availability of cultivable isolates and the requirement for biosafety level 3 or 4 (BSL3/4) containment for pathogenic hantaviruses (10). Here, we generated replication-competent recombinant vesicular stomatitis viruses (rVSVs) expressing an enhanced green fluorescent protein (eGFP) reporter and bearing codon-optimized Gn/Gc complexes from six ecologically and phylogenetically distinct hantaviruses: the HCPS-associated Black Creek Canal virus (BCCV, *Orthohantavirus nigrorivense*; carried by Hispid cotton rat) (27) and Laguna Negra virus (LANV, *Orthohantavirus mamorense*; small vesper mouse) (28); the Old World rodent-borne Dabieshan virus (DBSV, *Orthohantavirus dabieshanense*; Chinese white-bellied rat) (29) and Tigray virus (TIGV, *Orthohantavirus tigrayense*; Ethiopian white-footed mouse) (30, 31); the shrew-borne Cao Bang virus (CBNV, *Orthohantavirus caobangense*; Chinese mole shrew) (32) and Amga virus (MGAV, *Orthohantavirus artybashense*; Laxmann’s shrew) (33, 34). This panel includes viruses with distinct and incompletely defined zoonotic relevance. TIGV circulates in peridomestic rodents in Ethiopia, where orthohantavirus-reactive antibodies were detected in human sera (35). DBSV is genetically related to the prototypic HFRS-causing Hantaan virus (HTNV), whereas CBNV and MGAV are shrew-borne viruses with no established association with human disease (13, 29).

BCCV and LANV are particularly notable because authentic isolates exist for both, and each has been studied extensively in epidemiologic, reservoir, and animal-model settings. BCCV persists asymptomatically with horizontal transmission in its natural reservoir but is uniformly lethal in immunosuppressed Syrian hamsters (36–38), whereas LANV causes HCPS across South America and lethal pulmonary disease in Turkish hamsters (39). Despite this substantial body of work, the receptor requirements of both viruses remain undefined: BCCV entry was last examined before any hantavirus receptor was identified (40) and neither virus has been tested for PCDH1 dependence. To recover replication-competent rVSVs bearing Old World rodent-borne and shrew-borne glycoproteins, we engineered Gn/Gc constructs using previously established HTNV-based rescue-enhancing substitutions (23, 41). Here we show that all six Gn/Gc complexes mediate infection of human microvascular endothelial cells, consistent with systematic pseudotype screens showing that receptor-binding proteins from diverse animal viruses can function broadly in human cells (1). CRISPR/Cas9-based genetic depletion and cDNA complementation in human cells demonstrated that infection mediated by glycoproteins of BCCV and LANV—but not those of DBSV, TIGV, CBNV, or MGAV—depends on PCDH1. Concordantly, BCCV and LANV glycoproteins bound soluble PCDH1 with different efficiencies in sEC1-2 capture ELISAs, were competitively blocked at this interface by EC1-specific mAb-3305, and were neutralized by EC1-Fc and mAb-3305 in endothelial cell infection assays. Finally, ADI-42898 bound all six Gn/Gc-bearing particles and neutralized them with variable potency that correlated with naturally occurring amino acid variation at epitope sites corresponding to PUUV residues Q98, E750, and E768. Together, our findings expand the known diversity of hantavirus receptor usage and define human endothelial cell susceptibility and antibody sensitivity profiles across an expanded spectrum of hantavirus glycoproteins.

## Results

### Gn/Gc complexes from divergent rodent-and shrew-borne hantaviruses mediate infection of human endothelial cells

We evaluated whether Gn/Gc glycoproteins from six phylogenetically and ecologically distinct hantaviruses could mediate infection of human endothelial cells. The panel included the New World rodent-borne viruses Black Creek Canal virus (BCCV) and Laguna Negra virus (LANV), the Old World rodent-borne viruses Dabieshan virus (DBSV) and Tigray virus (TIGV), and the shrew-borne Cao Bang virus (CBNV) and Amga virus (MGAV) **(Fig. 1A).** Although cell culture isolates are available for BCCV and LANV (27, 28), authenticated isolates have not been reported for DBSV, TIGV, CBNV or MGAV, which are known from viral sequences recovered from reservoir tissues. To examine Gn/Gc-mediated entry under BSL2 conditions, we generated replication-competent recombinant vesicular stomatitis viruses (rVSVs) in which the native VSV G open reading frame was replaced with the Gn/Gc open reading frame from LANV, BCCV, DBSV, CBNV, TIGV, or MGAV. An enhanced green fluorescent protein (eGFP) reporter was included in the first transcriptional unit to permit quantitation of infected cells **(Fig. 1B, top)**. Consistent with established rVSV rescue strategies for Old World rodent-borne, mole-, and shrew-borne viruses, constructs from DBSV, CBNV, TIGV, and MGAV contained the Hantaan virus (HTNV)-derived rescue-enhancing substitutions I532K in the Gn cytoplasmic tail and S1094L in the membrane-proximal stem of Gc ectodomain (23, 41) **(Fig. 1B bottom)**. In selected constructs, adjacent residues were also replaced with their HTNV counterparts. These modifications promote plasma-membrane localization and rVSV incorporation of HTNV Gn/Gc and lie outside the receptor-binding interface and the mapped neutralizing-antibody epitopes evaluated here.

**Fig. 1.**
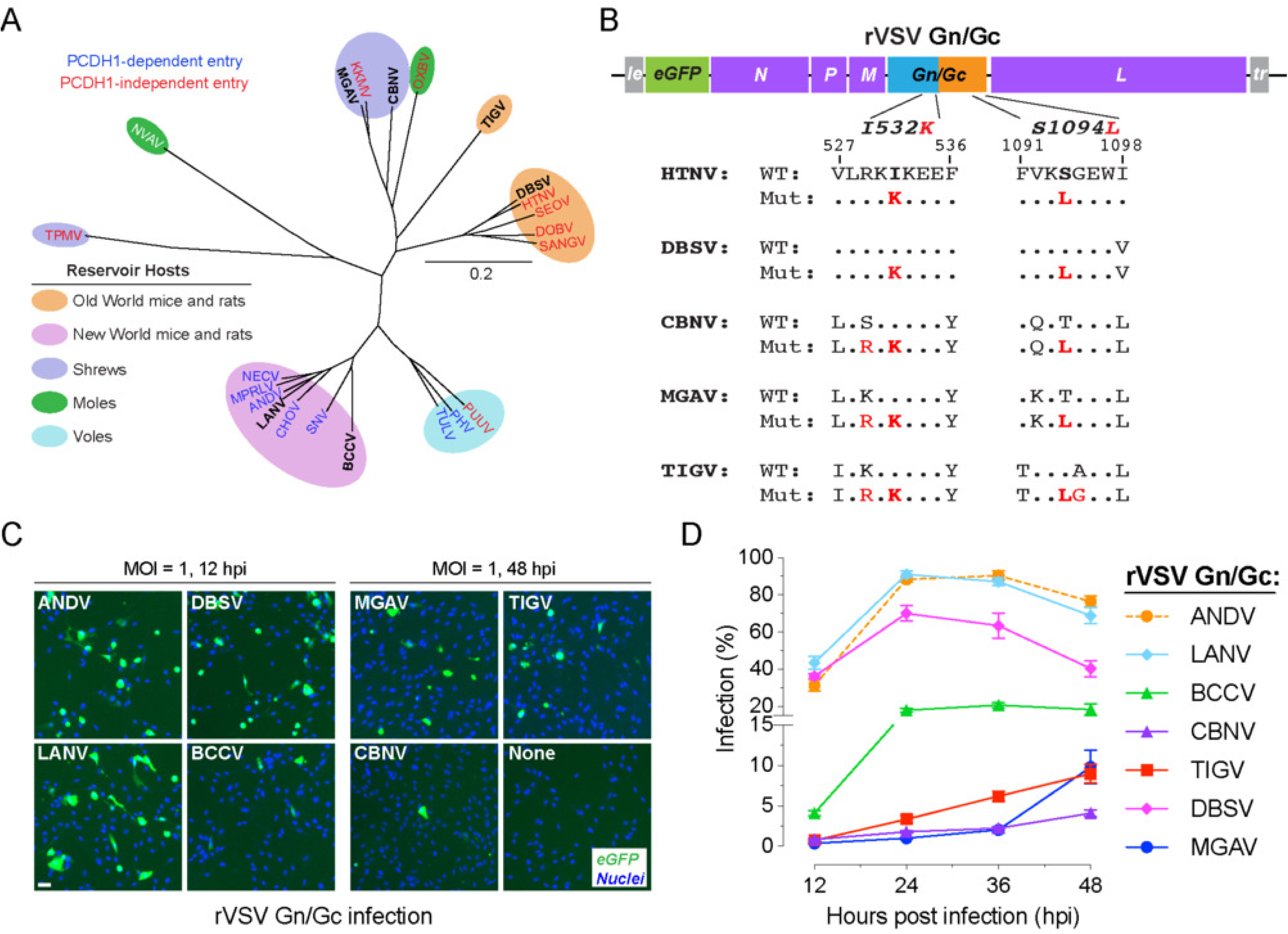
Divergent hantavirus Gn/Gc proteins support infection of human pulmonary microvascular endothelial cells with distinct kinetics. (A) Neighbor-joining phylogenetic tree showing the hantaviruses examined in this study together with related hantaviruses. The glycoprotein precursor amino acid sequences used for phylogenetic analysis were translated from GenBank nucleotide records for Andes virus (ANDV, AF291703.2), Hantaan virus (HTNV, M14627.1), Maporal virus (MPRLV, AY363179.1), Sin Nombre virus (SNV, L37903.1), Choclo virus (CHOV, DQ285047.1), Prospect Hill virus (PHV, X55129.1), Puumala virus (PUUV, M29979.1), Seoul virus (SEOV, S47716.1), Dobrava-Belgrade virus (DOBV, AJ410616.1), Necoclí virus (NECV, KF494345.1), Tula virus (TULV, Z69993.1), Nova virus (NVAV, KR072622.1), Oxbow virus (OXBV, FJ539167.1), Kenkeme virus (KKMV, KJ857337.1), Thottapalayam virus (TPMV, DQ825771.1), and Sangassou virus (SANGV, JQ082301.1). For the 6 viruses specific to this study, the GenBank accession numbers of the glycoprotein precursor amino acid sequences are Black Creek Canal virus (BCCV, L39950.1), Laguna Negra virus (LANV, AAB87603.1), Cao Bang virus (CBNV, ABR29829.1), Dabieshan virus (DBSV, AFG26309.1), Tigray virus (TIGV, AOR07001.1), and Amga virus (MGAV, AID54673.1). Colored ovals indicate their reservoir hosts. Hantaviruses previously reported to use PCDH1 for entry are indicated in blue, whereas viruses reported to enter independently of PCDH1 are indicated in red. White text denotes the partial PCDH1 dependence of Nova virus (NVAV). Viruses evaluated in this study are shown in bold. (B) Top, schematic of the recombinant vesicular stomatitis virus (rVSV) genome in which the VSV glycoprotein G gene is replaced by hantavirus Gn/Gc and an enhanced green fluorescent protein (eGFP) reporter is encoded in the first transcriptional unit. Bottom, alignment of Gn/Gc sequences showing rescue-enhancing amino acid substitutions (red) incorporated into the indicated rVSVs. Substitutions corresponding to I532K in the Gn cytoplasmic tail and S1094L in the Gc membrane-proximal stem are shown in bold. Dots indicate residues identical to those in the HTNV reference sequence. (C) Primary human pulmonary microvascular endothelial cells (HPMECs) were infected with the indicated rVSV Gn/Gc at an MOI of 1. Cells were imaged for eGFP expression and Hoechst 33342-stained nuclei at 12 h post-infection (hpi) for ANDV, LANV, BCCV, and DBSV, or at 48 hpi for MGAV, TIGV, and CBNV. Representative images from one of 3 independent experiments are shown. (D) HPMECs were infected with the indicated rVSV Gn/Gc at an MOI of 1, and infection was quantified over a 48 h time course. Data are shown as mean ± SEM; n = 9 from 3 independent experiments. Each *n* represents one well of a 96-well plate, with 500 to 1,000 cells scored per condition. Some error bars are smaller than the symbols.

Recovery of rVSV-BCCV required an S430C substitution. Because cysteine is highly conserved at the corresponding position among mammalian hantaviruses, the serine reported in the deposited BCCV GenBank sequence may reflect a sequencing error rather than authentic viral variation. Nanopore sequencing of rescued rVSV stocks identified no additional nonsynonymous changes beyond the engineered substitutions, except for G485R in rVSV-MGAV. This substitution has also been observed during rescue of rVSV bearing the related Kenkeme virus Gn/Gc (23) but not for most other hantavirus Gn/Gc proteins (20, 21, 42–44). G485, the terminal residue of the Gn ectodomain, is highly conserved among hantaviruses; however, the functional consequences of the G485R substitution remain unknown.

We then measured infection of primary human pulmonary microvascular endothelial cells (HPMECs) over 48 h at a multiplicity of infection (MOI) of 1 **(Fig. 1C and D)**. Virus inputs were normalized using titers determined in permissive Huh-7.5.1 cells. At 12 h postinfection (hpi), rVSV-LANV and rVSV-DBSV infection approached or exceeded that of the rVSV-ANDV control, with approximately 30-40% of cells becoming eGFP positive **(Fig. 1C and D)**. Infection by these viruses and by rVSV-ANDV peaked between 24 and 36 hpi, reaching approximately 70-90% eGFP positive cells **(Fig. 1D)**. In contrast, rVSV-BCCV infection was only minimally detectable at 12 hpi, at approximately 4%, and increased to a plateau of approximately 20% by 24 hpi **(Fig. 1D)**. rVSV-CBNV, rVSV-TIGV, and rVSV-MGAV produced slower and lower-level infection, becoming readily detectable only at 48 hpi, when approximately 4-8% of cells were eGFP positive **(Fig.1C and D).**

Thus, all six Gn/Gc complexes mediated infection of primary human endothelial cells but differed substantially in infection efficiency and kinetics. These data establish human endothelial-cell entry competence for LANV and BCCV Gn/Gc and provide functional evidence that the Gn/Gc complexes of DBSV, TIGV, CBNV, and MGAV can mediate cellular entry.

### BCCV and LANV Gn/Gc-mediated infection depends on PCDH1

We next tested whether the Gn/Gc complexes in this panel require PCDH1 for entry. Wild-type (WT), PCDH1-knockout cells transduced with empty vector (KO + EV), and PCDH1-knockout cells complemented with PCDH1 cDNA (KO + PCDH1) were infected with the indicated rVSVs. These genetic backgrounds in previously validated human osteosarcoma U2OS and telomerase-immortalized human microvascular endothelial (TIME) cell systems enable distinction between PCDH1-dependent and PCDH1-independent hantavirus entry pathways (20–23).

In U2OS cells, PCDH1 ablation reduced infection by rVSV-BCCV and rVSV-LANV by approximately 50-60%, similar to the reduction observed for the PCDH1-dependent rVSV-ANDV control **(Fig. 2A)**. PCDH1 re-expression restored rVSV-LANV and rVSV-ANDV infection. Infection by rVSV-BCCV also increased after PCDH1 complementation, reaching levels above those observed in WT cells. In contrast, PCDH1 status had little or no effect on infection by rVSV-DBSV, rVSV-CBNV, rVSV-TIGV, and rVSV-MGAV, paralleling the well-established PCDH1-independent infection of rVSV-HTNV.

**Fig. 2.**
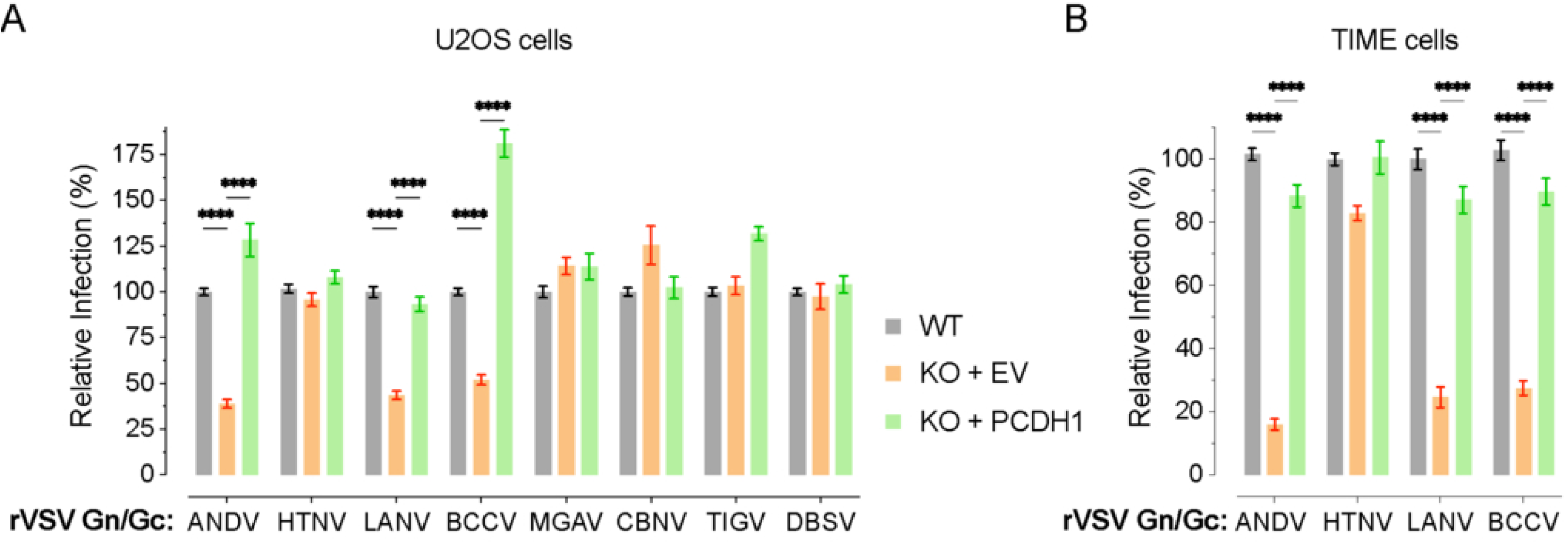
BCCV and LANV Gn/Gc-mediated infection is PCDH1-dependent in human U2OS and TIME cells. Wild-type (WT) cells, PCDH1-knockout (KO) cells complemented with empty vector (KO + EV), and PCDH1-KO cells complemented with PCDH1 cDNA (KO + PCDH1) were infected with the indicated rVSV Gn/Gc. Infection is presented relative to WT cells, which were set to 100%. Viruses were pre-titrated to estimate inocula for each rVSV that yielded 18–35% infection in WT U2OS and TIME cells. (A) Human osteosarcoma U2OS cells were analyzed 24 hpi. (B) Human endothelial TIME cells were analyzed 12 hpi. Data are shown as mean ± SEM; n = 8-9 from 3 independent experiments. Each n represents infection measured from 1,000 to 2,000 U2OS cells or 500 to 1,000 TIME cells per condition. Means within each rVSV group were compared by two-way ANOVA with Dunnett’s correction for multiple comparisons. ****, p < 0.0001.

We confirmed this phenotype in TIME cells, providing an endothelial cell context relevant to hantavirus tropism. In these cells, PCDH1 knockout reduced rVSV-BCCV and rVSV-LANV infection by approximately 75% relative to WT cells, comparable to the reduction observed for rVSV-ANDV **(Fig. 2B)**. rVSV-HTNV infection was not significantly affected by PCDH1 knockout. Re-expression of PCDH1 restored susceptibility to rVSV-BCCV, rVSV-LANV, and rVSV-ANDV toward WT levels.

The concordant loss-of-function and genetic complementation phenotypes in U2OS and endothelial cells identify PCDH1 as an entry factor for BCCV and LANV. By contrast, the absence of a comparable phenotype for DBSV, CBNV, TIGV, and MGAV indicates these divergent Gn/Gc complexes do not exhibit detectable dependence on PCDH1 in these cellular systems.

### PCDH1-targeting reagents inhibit BCCV and LANV Gn/Gc-mediated infection

Because BCCV and LANV Gn/Gc-mediated infection depended on PCDH1, we asked whether disrupting the PCDH1-Gn/Gc interaction could inhibit infection of endothelial cells. We tested EC1-Fc, a dimeric human Fc-tagged soluble EC1 decoy (23), against rVSV-BCCV and rVSV-LANV in HPMECs. As expected for a PCDH1-dependent control, EC1-Fc inhibited rVSV-ANDV infection in a concentration-dependent manner, reducing infection to approximately 5% of the no-decoy control at 300 nM **(Fig. 3A)**. rVSV-LANV was also inhibited in a concentration-dependent manner, although residual infection at 300 nM was greater than for rVSV-ANDV, at approximately 24%. EC1-Fc inhibited rVSV-BCCV less efficiently, with approximately 63% relative infection remaining at 300 nM. rVSV-HTNV infection was unaffected throughout the concentration range tested, supporting the specificity of EC1-Fc for PCDH1-dependent entry.

**Fig. 3.**
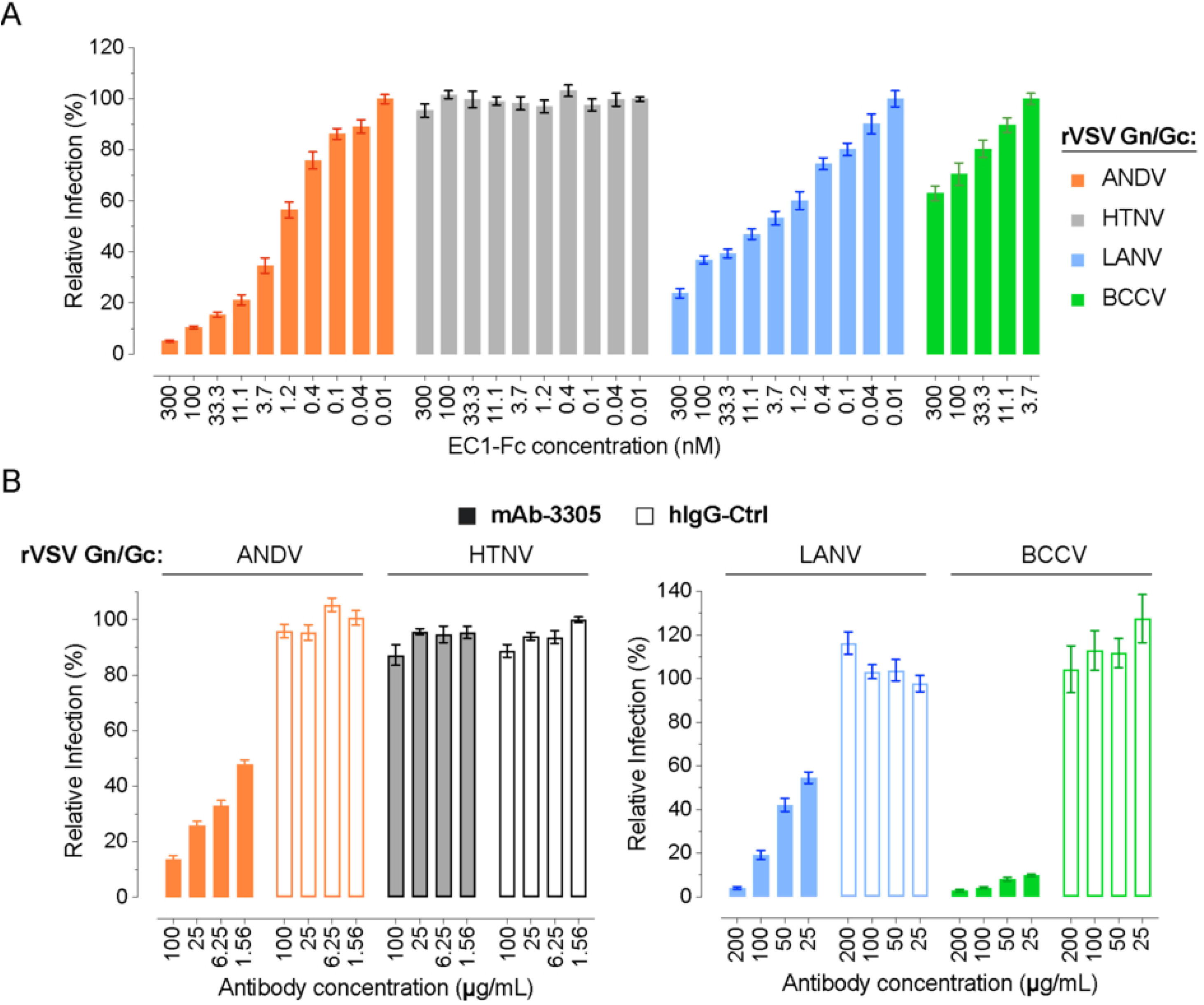
Soluble PCDH1 and PCDH1-targeting antibody inhibit BCCV and LANV Gn/Gc-mediated infection. Virus input was pretitrated to achieve approximately 18-30% infection in untreated control wells. (A) HPMECs were infected with the indicated rVSV Gn/Gc after preincubation with serial dilutions of soluble PCDH1 EC1-Fc. At 8 hpi, infection was quantified relative to the no-EC1-Fc control, which was set to 100%. (B) HPMECs were pretreated with the indicated concentrations of the EC1-specific anti-PCDH1 monoclonal antibody mAb-3305 or a nonspecific human IgG control (hIgG-Ctrl) before infection with the indicated rVSV Gn/Gc. At 8 hpi, infection was quantified relative to the no-antibody control, which was set to 100%. Data are shown as mean ± SEM; n = 6-9 from 3 independent experiments. Each *n* represents the infection status of 500 to 1,000 cells per condition.

We also evaluated mAb-3305, an EC1-specific human monoclonal antibody that blocks PCDH1-dependent hantavirus entry (21). HPMECs were pretreated with mAb-3305 or a nonspecific human IgG control before infection. mAb-3305 inhibited rVSV-LANV, rVSV-BCCV, and rVSV-ANDV infection in a concentration-dependent manner, whereas rVSV-HTNV infection remained unaffected **(Fig. 3B)**. Notably, rVSV-BCCV was particularly sensitive to mAb-3305, with infection reduced to approximately 2-10% of control levels across the concentrations tested. By comparison, residual infection ranged from approximately 4-55% for rVSV-LANV and approximately 14-48% for rVSV-ANDV. The nonspecific human IgG control had negligible effect on infection.

Both PCDH1-directed interventions therefore inhibited BCCV and LANV Gn/Gc-mediated infection, consistent with a requirement for EC1 in viral entry, but their relative effects differed between the two viruses. BCCV was less susceptible to the soluble EC1-Fc decoy but more sensitive to cell-surface PCDH1 blockade by mAb-3305. These distinct inhibition profiles are consistent with differences in the apparent PCDH1 engagement of BCCV and LANV Gn/Gc, which we investigated further using PCDH1 capture assays.

### BCCV and LANV Gn/Gc-bearing particles engage PCDH1 with different apparent efficiencies

We used a previously established soluble PCDH1-EC1-2 capture ELISA to assess PCDH1 engagement by BCCV and LANV Gn/Gc-bearing particles (20–23). rVSV inputs were first normalized to VSV matrix protein (VSV M) abundance in pelleted virions, thereby controlling for relative particle input **(Fig. 4A, top)**. Equivalent amounts of biotinylated rVSVs were then added to sEC1-2-coated ELISA plates, and captured particles were detected by streptavidin-HRP. rVSV-ANDV or rVSV-LANV produced comparable dose-dependent capture signals **(Fig. 4A, bottom)**. rVSV-BCCV was also captured in a dose-dependent manner, but the signal was lower than that produced by rVSV-ANDV or rVSV-LANV at corresponding virus inputs. The PCDH1-independent rVSV-HTNV control showed minimal capture. These findings support specific engagement of soluble PCDH1 by both BCCV and LANV Gn/Gc-bearing particles, with a lower apparent PCDH1-capture efficiency for BCCV under these assay conditions.

**Fig. 4.**
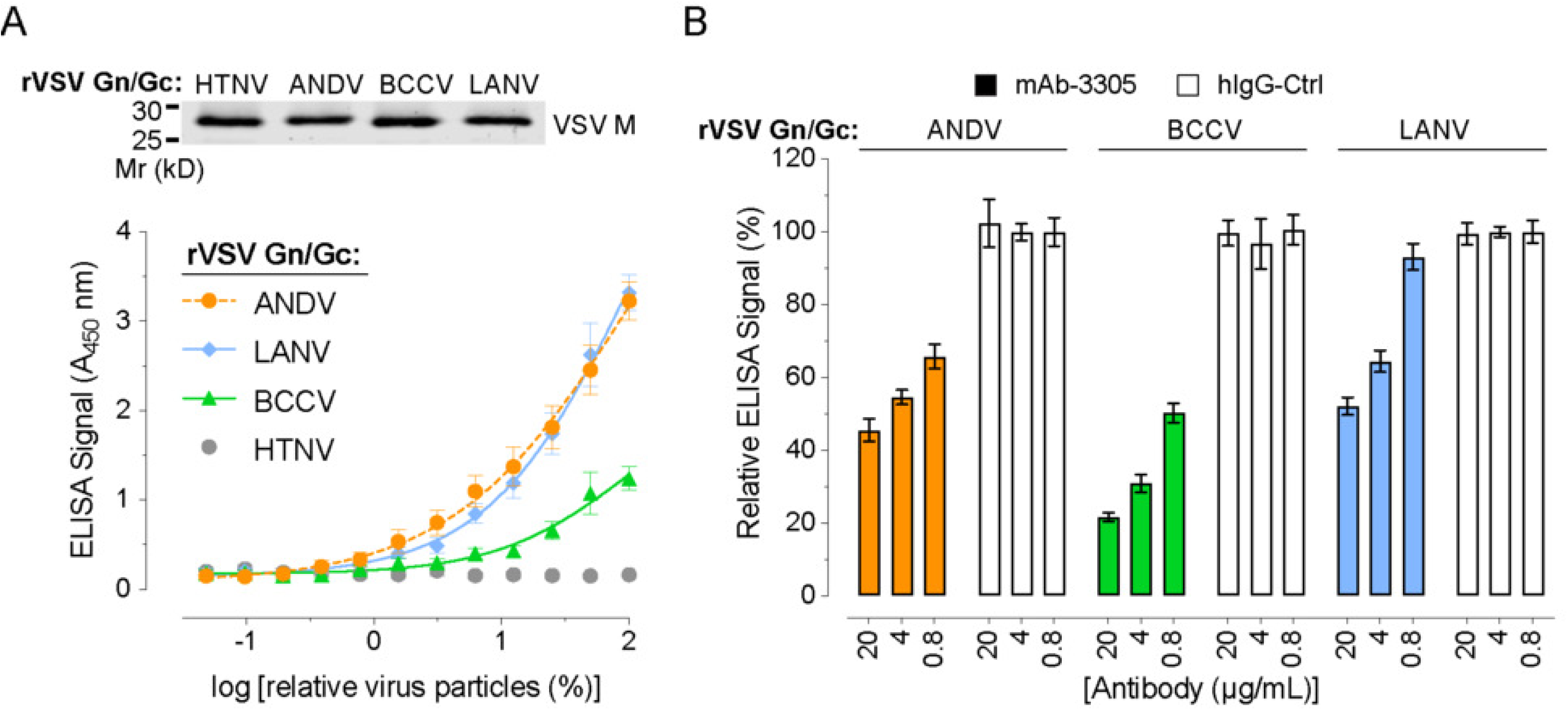
BCCV and LANV Gn/Gc bind PCDH1 with distinct efficiencies. (A) Top, representative immunoblot of VSV matrix protein (VSV M) showing normalization of pelleted rVSV particles bearing the indicated Gn/Gc proteins. Bottom, normalized amounts of biotinylated rVSVs were serially diluted and applied to ELISA plates coated with soluble PCDH1 extracellular domain 1 and 2 (sEC1-2). Captured virus was quantified by ELISA at 450 nm. Data are shown as mean ± SEM; n = 9 from 3 independent experiments. (B) PCDH1-dependent rVSV capture was assessed by preincubating sEC1-2-coated ELISA plates with mAb-3305 or a nonspecific human IgG control (hIgG-Ctrl) before addition of the indicated normalized rVSVs. ELISA signal is shown relative to the corresponding no-antibody control set to 100%, which corresponds to an OD of 1.2 to 1.5. Data are shown as mean ± SEM; n = 7-9 from 3-4 independent experiments. Some error bars are smaller than the symbols. Mr denotes relative molecular weight.

To confirm the specificity of this interaction, we performed competition ELISA using the EC1-targeting mAb-3305 (20–23). Preincubation of sEC1-2 with mAb-3305 reduced capture of rVSV-LANV and rVSV-BCCV in a concentration-dependent manner, whereas the nonspecific human IgG control had no measurable effect **(Fig. 4B)**. Together with the genetic-dependence and infection-blockade experiments, these capture data support EC1-mediated PCDH1 engagement by both BCCV and LANV Gn/Gc. The lower BCCV capture signal, however, could reflect reduced incorporation of BCCV Gn/Gc into rVSV particles, a difference in PCDH1 engagement per incorporated glycoprotein complex, or both.

### Reduced BCCV Gn/Gc incorporation contributes to, but does not fully explain, weaker PCDH1 capture

The capture ELISAs in **Fig. 4** used rVSV inputs normalized to VSV M abundance, which controls for particle input but not necessarily for the amount of incorporated Gn/Gc into each particle. We therefore compared Gn/Gc incorporation among pelleted rVSV-ANDV, rVSV-LANV, and rVSV-BCCV preparations. Rabbit antiserum raised against an ANDV Gc peptide recognized Gc from all three viruses by immunoblotting, consistent with the fact that the immunizing peptide is identical in LANV Gc and differs by only two residues in BCCV Gc **(Fig. 5A and B)**. When equivalent VSV M-normalized rVSV inputs were compared, the BCCV Gc signal was lower than the ANDV and LANV Gc signals **(Fig. 5B)**. Quantification of the Gc/VSV M ratio confirmed that rVSV-BCCV contained significantly less Gc relative to VSV M than rVSV-ANDV or rVSV-LANV **(Fig. 5C)**. Silver staining provided an independent assessment of virion-associated proteins and likewise showed less prominent Gn/Gc bands for rVSV-BCCV preparations than in rVSV-ANDV or rVSV-LANV preparations **(Fig. 5D)**. These results indicate that BCCV Gn/Gc is incorporated into rVSV particles less efficiently than ANDV or LANV Gn/Gc.

**Fig. 5.**
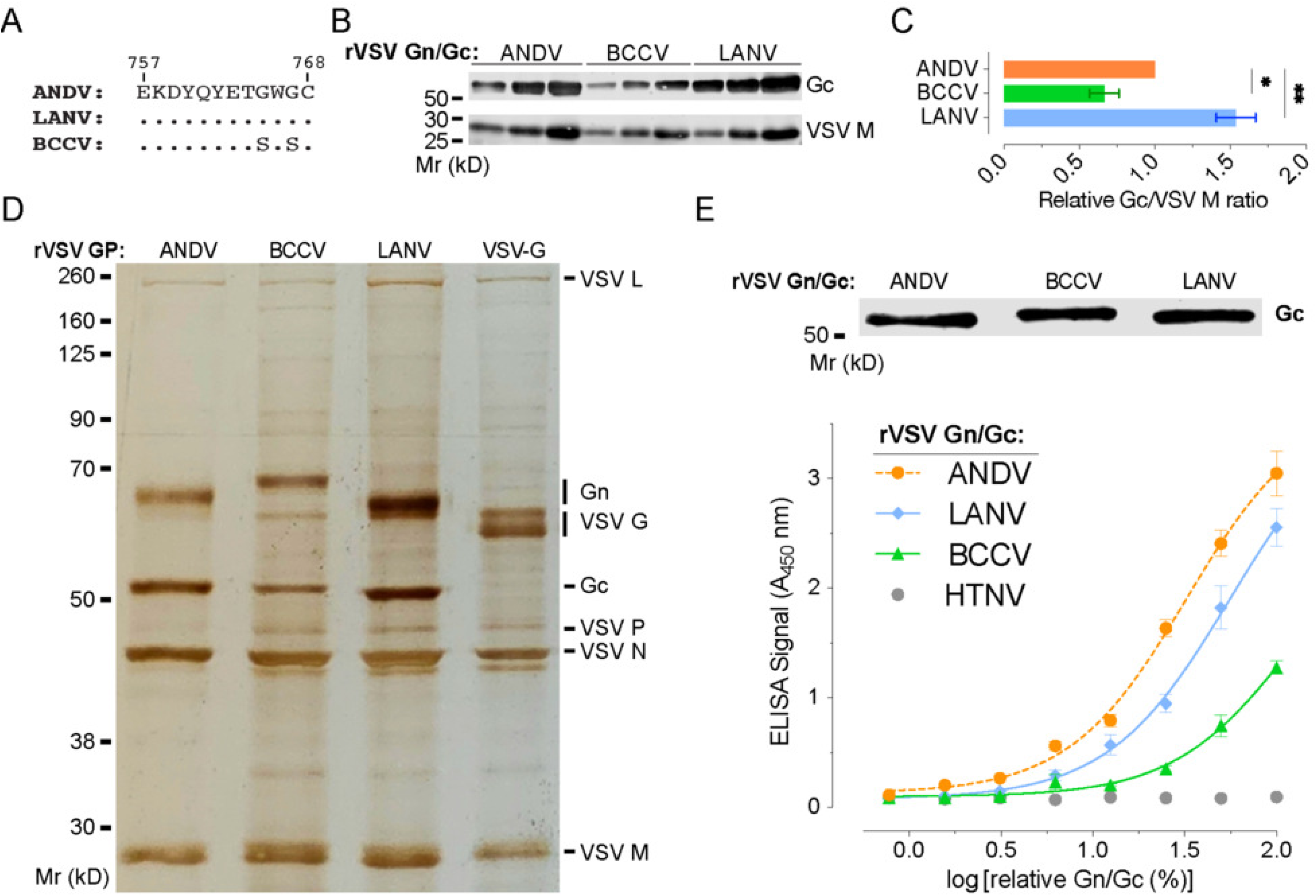
Reduced BCCV Gn/Gc incorporation and lower PCDH1 binding contribute to diminished BCCV capture. (A) Alignment of amino acid sequence from the indicated hantavirus Gc proteins corresponding to ANDV Gc residues 757-768, the peptide used to generate the anti-Gc antiserum used in panel B. Dots indicate amino acids identical to the ANDV sequence. (B) Representative immunoblots of Gc and VSV M proteins in increasing volumes of pelleted rVSVs bearing ANDV, BCCV or LANV Gn/Gc. (C) Quantification of the relative Gc/VSV M ratio from the immunoblot analysis in panel B. Data are shown as mean ± SEM; n = 9 from 3 independent experiments. Two-way ANOVA with Dunnett’s correction for multiple comparisons. *, p < 0.05, **, p < 0.01. (D) Representative silver-stained SDS-PAGE gel of pelleted rVSV particles normalized by VSV M content. One representative gel from at least 3 independent experiments is shown. (E) Top, representative immunoblot showing normalized Gc inputs from rVSV particles bearing the indicated Gn/Gc proteins. Bottom, serial twofold dilutions of rVSV particles normalized by Gc content were applied to sEC1-2-coated ELISA plates, and captured virus was quantified by ELISA at 450 nm. Data are shown as mean ± SEM; n = 6 from 3 independent experiments. Mr denotes relative molecular weight. Some error bars are smaller than the symbols.

We next asked whether reduced glycoprotein incorporation fully accounted for the weaker BCCV PCDH1-capture signal. rVSV inputs were adjusted according to relative Gc abundance before addition to sEC1-2-coated plates **(Fig. 5E, top).** Even after this normalization, rVSV-BCCV was captured less efficiently than rVSV-ANDV and rVSV-LANV across the range of inputs tested, whereas rVSV-HTNV remained near background **(Fig. 5E, bottom).** Thus, reduced BCCV Gn/Gc incorporation contributes to the lower PCDH1-capture signal but does not fully explain it.

### ADI-42898 neutralizes rVSVs bearing divergent hantavirus Gn/Gc with variable potency

Having established that BCCV and LANV use PCDH1 for entry, and that targeting this interaction is effective at reducing their infection, we next evaluated a receptor-independent, spike-directed neutralization strategy. ADI-42898 is a human broadly neutralizing antibody isolated from a PUUV-experienced donor that recognizes a quaternary epitope spanning the Gn capping loop and Gc fusion loops (24, 26). Comparison of the mapped ADI-42898 epitope across the viruses in our panel showed substantial amino acid variation at positions corresponding to PUUV Gn/Gc residues: Q98, E750, and E768 **(Fig. 6A)**. We first measured relative reactivity of ADI-42898 with rVSV-associated Gn/Gc proteins by ELISA using VSV M-normalized particle inputs **(Fig. 6B)**. ADI-42898 reacted most strongly with rVSVs bearing DBSV, LANV and MGAV Gn/Gc **(Fig. 6C).** Signals for rVSVs-ANDV, rVSV-BCCV, rVSV-CBNV, or rVSV-TIGV were lower but remained higher than the VSV-G-bearing particles. Thus, despite marked sequence variation in the mapped epitope, ADI-42898 retained recognition of Gn/Gc complexes from all six viruses in the panel.

**Fig. 6.**
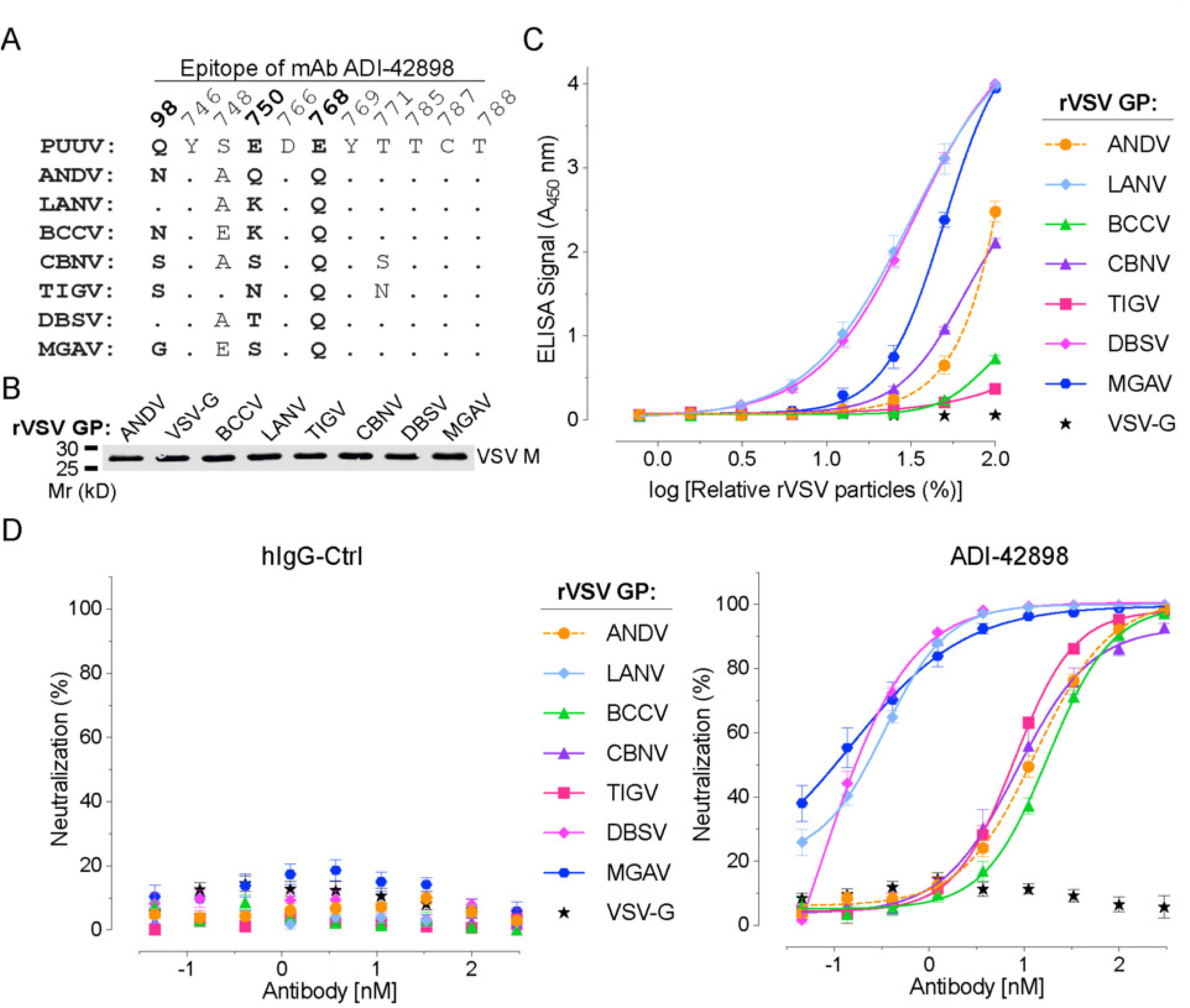
Broadly neutralizing human mAb ADI-42898 recognizes and neutralizes rVSVs bearing divergent hantavirus Gn/Gc proteins. (A) Alignment of amino acid sequences within the mapped ADI-42898 epitope in the Gn/Gc proteins of the indicated hantaviruses. Residues are numbered according to the PUUV Gn/Gc sequence, and dots indicate amino acids identical to PUUV. (B) Representative VSV M immunoblot showing normalization of pelleted rVSV particles used in the ADI-42898 binding assay. (C) Serial twofold dilutions of normalized rVSV particles applied to ELISA plates and probed with ADI-42898. Antibody binding was measured as absorbance at 450 nm. (D) The indicated rVSVs were preincubated with serial dilutions of unrelated human IgG control or ADI-42898 before infection of Vero cells; BCCV was tested in HPMECs. Infection was quantified relative to the no-antibody control, which was set to 100%. VSV carrying authentic VSV G served as a negative control in the ELISA and neutralization assays. Data are shown as mean ± SEM; n = 8-12 from 3-4 independent experiments. Each *n* in panel D represents the infection status of 500 to 1,000 cells per condition. Some error bars are smaller than the symbols. Mr denotes relative molecular weight.

We next determined whether the differences in ELISA reactivity were reflected in neutralization potency. Neutralization assays were performed in Vero cells, except for rVSV-BCCV, which was evaluated in HPMECs because of low infectivity in Vero cells. Preincubation with ADI-42898, but not the negative control (a non-specific human IgG antibody) **(Fig. 6D, left),** reduced infection by every rVSV tested **(Fig. 6D, right)**.

However, the antibody concentration required for neutralization varied among viruses **(Fig. 6D, right)**. rVSVs bearing ANDV, BCCV, CBNV, or TIGV Gn/Gc required higher concentrations of ADI-42898 to achieve 50% neutralization, with IC_50_ values ranging from 10 to 20.32 nM. In contrast, rVSVs bearing DBSV, LANV, and MGAV Gn/Gc were more sensitive, with IC_50_ values ranging from 0.13 to 0.27 nM. These findings show that ADI-42898 can recognize and neutralize infection mediated by Gn/Gc complexes from phylogenetically divergent rodent-and shrew-borne hantaviruses. Stronger particle-associated ELISA reactivity for DBSV, LANV, and MGAV broadly paralleled their greater sensitivity to neutralization. Taken together, these data indicate that sequence variation within the ADI-42898 epitope is compatible with broad recognition, while virus-specific differences in Gn/Gc reactivity are associated with substantial variation in neutralization potency.

## Discussion

Our comparative analysis of six divergent hantavirus Gn/Gc complexes identified human endothelial cell entry competence, PCDH1 usage, and sensitivity to entry-directed countermeasures as separable phenotypes. All six Gn/Gc-bearing rVSVs infected primary human pulmonary microvascular endothelial cells, although their kinetics and extent of infection differed substantially **(Fig. 1)**. rVSV-LANV and rVSV-DBSV infected HPMECs rapidly and efficiently, whereas rVSV-BCCV showed lower-level infection, and rVSV-CBNV, rVSV-TIGV, and rVSV-MGAV produced delayed, limited infection. These differences indicate that endothelial cell infection mediated by hantavirus glycoproteins is quantitatively heterogeneous and may reflect variation in attachment, receptor use, internalization, fusion, or glycoprotein-dependent spread.

However, entry in this rVSV system should not be equated with authentic-virus zoonotic potential or pathogenicity, which additionally depend on post-entry replication, immune evasion, host ecology, and the capacity to produce endothelial dysfunction. This distinction between rVSV-mediated entry and authentic-virus zoonotic potential or pathogenicity is particularly important for DBSV, CBNV, TIGV, and MGAV, which remain incompletely characterized and lack the epidemiologic and experimental foundation available for LANV and BCCV.

LANV and BCCV were prioritized here because both are medically important New World rodent-borne hantaviruses for which the molecular determinants of entry have remained poorly defined. LANV was initially associated with a large HCPS outbreak in the Paraguayan Chaco, where viral sequences from patients and *Calomys laucha* rodents established a strong reservoir-human disease association; LANV and LANV-like viruses have subsequently been associated with HCPS cases and *Calomys* hosts in Bolivia, Argentina, and Brazil (28, 45, 46). LANV can also produce pulmonary disease in immunocompetent Turkish hamsters, further supporting its pathogenic potential(39). BCCV was isolated from hispid cotton rats (*Sigmodon hispidus*) in southern Florida after recognition of a severe HCPS-like illness in a human patient. Subsequent studies have examined its ecology in reservoir hosts, transmission, and interaction with host cells (27, 36, 37, 47). BCCV also caused uniformly lethal pulmonary disease in immunosuppressed Syrian hamsters, providing additional experimental evidence of pathogenic potential in a heterologous mammalian host (38). Together with its prior association with severe HCPS in Florida, these findings support BCCV as a medically important hantavirus whose molecular entry determinants warrant further investigation.

Our genetic and biochemical experiments establish PCDH1 as a functional entry factor for both LANV and BCCV. PCDH1 knockout reduced LANV and BCCV infection in U2OS and TIME cells, with a more pronounced phenotype in endothelial cells, and PCDH1 cDNA complementation restored susceptibility **(Fig. 2)**. In parallel, both viruses showed EC1-dependent capture in the soluble PCDH1 assay, and binding was inhibited by the EC1-directed antibody mAb-3305 **(Fig. 4)**. These convergent findings identify BCCV and LANV as additional PCDH1-using hantaviruses and expand the set of HCPS-associated viruses with experimentally demonstrated PCDH1 dependence beyond ANDV and SNV. PCDH1 usage was previously reported for the New World viruses Maporal virus (MPRLV), Prospect Hill virus (PHV), and Choclo virus (CHOV) and, more recently, for Necoclí virus (NECV) and phylogenetically distinct Tula virus (TULV), with a weaker phenotype observed for Nova virus (NVAV) (20, 21, 23). The present work is distinct in defining PCDH1 dependence for two epidemiologically important HCPS-associated viruses and in revealing differences in their rVSV particle-associated glycoprotein incorporation, apparent PCDH1 engagement, and sensitivity to receptor-directed inhibitors.

Although both BCCV and LANV use PCDH1, their interactions with the receptor are not equivalent. When rVSV inputs were normalized to VSV M protein, rVSV-BCCV particles showed lower EC1 capture than rVSV-LANV or rVSV-ANDV particles **(Fig. 4)**. BCCV Gn/Gc was also incorporated less efficiently into rVSV particles **(Fig. 5A-D)**, initially suggesting that lower spike abundance contributed to reduced capture. However, adjustment of particle inputs according to Gc abundance did not eliminate this difference **(Fig. 5E)**, indicating that reduced Gn/Gc incorporation is not the sole explanation. Thus, reduced Gn/Gc incorporation is not the sole explanation for weaker BCCV capture. The most conservative interpretation is that BCCV displays lower apparent PCDH1 engagement than LANV and ANDV under the conditions of the capture assay. This result should not be interpreted as a direct measurement of affinity, because virion-capture assays can be influenced by glycoprotein density, multivalent avidity, epitope accessibility, conformational heterogeneity, and particle orientation. Nevertheless, the data support a meaningful functional distinction in how BCCV and LANV Gn/Gc engage PCDH1.

These differences in apparent receptor engagement were accompanied by distinct responses to PCDH1-directed inhibitors. LANV was more sensitive than BCCV to the soluble EC1-Fc decoy, whereas BCCV was particularly sensitive to the EC1-blocking mAb-3305 **(Fig. 3)**. These results show that the relative sensitivity of PCDH1-dependent hantaviruses can differ between soluble EC1 decoys and PCDH1-blocking antibodies, a pattern also observed for other PCDH1-dependent hantaviruses (21, 23). In the EC1-Fc assay, virions were preincubated with soluble decoy before encountering cells, whereas in the mAb-3305 assay, cell-surface PCDH1 was preoccupied before virus addition. The BCCV phenotype is consistent with a pattern observed previously for SNV and, more recently, NECV; viruses with weaker apparent PCDH1 binding were less sensitive to EC1-containing soluble decoys but more sensitive to mAb-3305-mediated receptor blockade (21, 23). This recurrent pattern suggests that the relative activity of these interventions may depend on the efficiency or mode of Gn/Gc engagement with PCDH1. However, the distinct inhibition profiles may also reflect experimental configuration, inhibitor stoichiometry, avidity, receptor-binding geometry, or accessibility of the EC1-binding interface. Direct biophysical measurements and matched competition experiments will be required to distinguish these nonexclusive mechanisms. Finally, ADI-42898 provided a potent receptor-independent strategy across the six viruses examined here: it bound and neutralized every Gn/Gc-bearing rVSV, with IC_50_ values ranging from approximately 0.13 to 20.32 nM **(Fig. 6).** rVSVs bearing DBSV, LANV and MGAV glycoproteins were most sensitive, whereas those bearing ANDV, BCCV, CBNV, or TIGV glycoproteins required higher antibody concentrations for equivalent neutralization. Nevertheless, all six panel viruses were susceptible at low-nanomolar to low-double-digit nanomolar IC_50_ concentrations. ADI-42898 was originally isolated from a PUUV-experienced individual and recognizes a prefusion quaternary Gn/Gc epitope involving the Gn capping region and conserved features surrounding the Gc fusion loops (24, 26). Unlike the many hantavirus neutralization antibodies that are restricted to individual viruses or closely related clades, antibodies targeting conserved Gc-associated or quaternary Gn/Gc surfaces can achieve substantially broader activity (24, 25). The broader panel evaluated by Word et al. further indicates that ADI-42898 activity is virus dependent across divergent hantavirus Gn/Gc complexes and does not extend equally to every rVSV tested (23). Nevertheless, its potent activity against all six viruses examined here, together with previously reported neutralization and protection against multiple HCPS-and HFRS-associated hantaviruses, supports ADI-42898 as a promising broadly active candidate against pathogenic hantaviruses (24, 26). Stronger particle-associated ADI-42898 reactivity and greater neutralization sensitivity for rVSV-DBSV, rVSV-LANV, and rVSV-MGAV suggest that virus-specific epitope presentation contributes to the observed potency differences. Sequence differences at positions corresponding to PUUV Q98, E750, and E768 are candidate determinants, but mutagenesis and structural analyses will be required to establish their contribution to sensitivity. Together, these findings define PCDH1-dependent entry by LANV and BCCV, reveal virus-specific differences in receptor engagement and receptor-directed inhibition, and identify conserved Gn/Gc surfaces as complementary targets for broadly reactive hantavirus countermeasures.

## Materials and methods

### Cells

Human embryonic kidney 293FT, human hepatoma Huh-7.5.1 [a generous gift from Charles M Rice (48) and Francis V Chisari (49)], and African green monkey kidney Vero cells were maintained in high-glucose Dulbecco’s modified Eagle mediaum (DMEM, Cytiva) supplemented with 10% fetal bovine serum (FBS; GeminiBio), except for Vero cells, which received 2% FBS. Human osteosarcoma U2OS cells were maintained in McCoy’s 5A medium (Corning) supplemented with the 10% FBS. All these media also contained 1% GlutaMAX (Gibco), and 1% penicillin-streptomycin (Gibco). Primary human pulmonary microvascular endothelial cells (HPMECs, Lonza) and telomerase-immortalized human microvascular endothelial (TIME) cells were maintained in VascuLife endothelial medium with the manufacturer’s EnGS supplement kit (Lifeline Cell Technology), as previously described (23). Suspension ExpiCHO-S (Thermo Fisher) and Freestyle 293-F (Thermo Fisher) cells used for protein and antibody expression were maintained in non-baffled shaker flasks in their respectively commercial expression media (Gibco) supplemented with 1% penicillin-streptomycin (Gibco) and 1% GlutaMAX (Gibco). Adherent cultures were maintained at 37℃, 5% CO2; suspension cultures were maintained at 37℃, 8% CO2, shaking at 125-130 rpm. All cells were tested monthly for mycoplasma and were negative.

### PCDH1-knockout and PCDH1-complemented cell lines

PCDH1-knockout U2OS and TIME cells, and their corresponding PCDH1 cDNA-complemented derivatives, were generated and validated previously by CRISPR/Cas9-mediated genome editing (20, 21) and were kindly provided by Dr. Kartik Chandran (Albert Einstein College of Medicine). Complemented U2OS cells were maintained under puromycin selection (2 μg/mL) as described previously (21).

### Plasmids

Codon-optimized Gn/Gc coding sequences for BCCV (GenBank L39950.1), LANV (strain 510B, AAB87603.1), CBNV (strain 3, ABR29829.1), DBSV (strain Yongjia-Nc-58, AFG26309.1), TIGV (strain ET2121, AOR07001.1), and MGAV (strain AH301, AID54673.1) were synthesized by Twist Biosciences and cloned in place of the native glycoprotein gene within a T7-driven recombinant VSV genomic plasmid encoding eGFP, using the same overall cloning strategy described previously (23, 41, 42, 50, 51). To enable rescue of the Old World rodent-borne and shrew-borne constructs, HTNV-derived rescue-enhancing substitutions (I532K in the Gn cytoplasmic tail and S1094L in the Gn membrane-proximal stem), together with selected adjacent HTNV-matched residues **(Fig. 1B)**, were introduced into DBSV, CBNV, TIGV and MGAV Gn/Gc sequences by standard molecular cloning, following the general substitution strategy validated for Old World hantavirus Gn/Gc rescue(23, 41).

The plasmids for the expression of sEC1-2, human Fc-tagged PCDH1 EC1 (EC1-Fc), heavy and light chains of mAb-3305 and ADI-42898 were described previously (21, 23).

### Generation of recombinant VSVs bearing hantavirus glycoproteins

rVSVs bearing LANV, BCCV, CBNV, DBSV, TIGV, or MGAV Gn/Gc were rescued in 293FT cells using a standard VSV plasmid-based rescue system, as previously described (23, 41, 42, 50, 51). Genome plasmids encoding Gn/Gc were co-transfected with helper plasmids encoding VSV N, P, M, G, L and T7 RNA polymerase. Rescued virus was amplified in Huh-7.5.1 cells. rVSV-ANDV and rVSV-HTNV were a generous gift from Kartik Chandran, and their generation has been described previously (42).

Glycoprotein identity and sequence was confirmed for every stock by RT-PCR amplification of the Gn/Gc-coding region from viral genomic RNA followed by Nanopore sequencing (Plasmidsaurus). Infectivity was quantified by automated enumeration of eGFP-positive cells using a Cytation 5 imager (Agilent Technologies) with its onboard Gen5 software essentially as described previously(23). For ELISA, immunoblotting and silver-staining experiments, rVSV particles were concentrated by ultracentrifugation over a sucrose cushion using previously described methods (23).

### Expression and purification of sE1-2, EC1-Fc and monoclonal antibodies

Soluble PCDH1 EC1-2 (sEC1-2) was expressed in Freestyle 293-F cells and purified using nickel-affinity chromatography exactly following the published methods(21). EC1-Fc, mAb-3305, and ADI-42898 were expressed by transient transfection of ExpiCHO-S cells using ExpiFectamine CHO reagent according to the manufacturer’s protocol as described earlier (23). Heavy and light-chain plasmids were co-transfected in equal amounts for the antibodies. Culture supernatants were harvested 7-14 days post-transfection, clarified by centrifugation, and incubated with Protein A agarose to capture the secreted protein. Bound protein was washed and eluted using Pierce binding and elution buffers (Thermo Fisher), buffer-exchanged into HEPES-NaCl (pH 7.4) using PD-10 columns (Cytiva) and concentrated using 10 (EC1-Fc) or 30-kDa (mAbs) MWCO Amicon filters (Sigma-Aldrich). Protein concentration was determined by absorbance at 280 nm.

### sEC1-2 capture and competition ELISAs

Capture ELISAs were performed largely using previously described methods (21–23). rVSV-ANDV and rVSV-HTNV were used as positive and negative controls, respectively. VSV M-normalized, FSL-biotin-labeled rVSV particles bearing BCCV or LANV Gn/Gc were added in serial dilution to high-binding plates coated with sEC1-2 (100 ng/well).

Bound virus was detected with streptavidin-HRP (Thermo Fisher) and 1-step Ultra TMB substrate (Thermo Fisher), and absorbance was read at 450 nm (Cytation 5, Agilent Technologies). ELISA data were fit to a four-parameter logistic (4PL) curve with a variable slope in GraphPad Prism v11, using log-transformed concentrations as x values. For competition ELISA, sEC1-2-coated plates were preincubated with serially diluted mAb-3305 or a nonspecific human IgG control before addition of a fixed amount of rVSV, pre-normalized to obtain an OD of 1.2-1.5 in the absence of antibody. Competition ELISA signals were normalized to the no-antibody control, defined as 100%, and expressed as relative signal (%).

### ADI-42898 binding ELISA

Reactivity of ADI-42898 with particle-associated Gn/Gc was assessed as described previously (23, 24), using rVSV-ANDV and VSV-G-bearing particles as positive and negative controls, respectively. VSV M-normalized rVSV particles bearing BCCV, LANV, CBNV, DBSV, TIGV, or MGAV Gn/Gc were serially diluted and used to coat ELISA plates overnight. Next day, it was followed by blocking, incubation with ADI-42898 (1 µg/mL), and detection with HRP-conjugated anti-human IgG (Thermo Fisher) and TMB substrate (Thermo Fisher), with absorbance read at 450 nm.

### Gn/Gc incorporation: immunoblotting and silver staining

These assays were performed using previously described methods (23). VSV M-normalized, pelleted rVSV particles bearing ANDV, LANV, or BCCV Gn/Gc were denatured and resolved by SDS-PAGE. Gc was detected by immunoblotting using rabbit antisera raised against an ANDV Gc peptide (**Fig. 5A**, Pacific Immunology). VSV M was detected in parallel using mouse anti-VSV-M mAb 23H12. Band intensities were quantified an Odyssey CLx Infrared Imaging System (LI-COR). For an unbiased assessment of total virion-associated protein, parallel gels were subjected to silver staining (52) and visualized using an I2R Glow Box transilluminator. VSV-G-bearing particles were used as an additional control for silver staining.

### Neutralization assays

Neutralization assays and data acquisition were performed using our previously published methods (23). For mAb-3305 neutralization, HPMECs were preincubated for 1 h at 37℃ with serially diluted mAb-3305 or a nonspecific human IgG control before infection with a pretitrated rVSV dose yielding approximately 18-30% baseline infection; infection was scored at 8 h post-infection (hpi). For EC1-Fc and ADI-42898 neutralization, rVSV particles were preincubated with serially diluted EC1-Fc, ADI-42898 or a nonspecific human IgG control for 1 h at 37℃ before addition to cells for an 8 h incubation period. All EC1-Fc assays were performed on HPMECs. ADI-42898 and IgG control assays were performed on Vero cells for all rVSVs except rVSV-BCCV, which was tested on HPMECs. In all assays, cells were fixed with 3% formaldehyde, nuclei were stained with Hoechst 33342, and infection was quantified by automated eGFP-positive cell counting (Cytation 5, Agilent Technologies) and normalized to no-antibody/no-decoy control.

### Endothelial cell infection kinetics

rVSV titers were determined on Huh-7.5.1 cells at 8 hpi. HPMECs were then infected at an MOI of 1 with each rVSV, and infection was scored every 12 h through 48 h post-infection using the fixation, staining, and imaging procedures described above.

### PCDH1-dependence infection assays

WT, PCDH1 knockout cells complemented with empty vector (KO + EV) or PCDH1 cDNA (KO + PCDH1) U2OS and TIME cells were infected in triplicate with a pretitrated rVSV dose yielding 18-35% infection in WT cells, infection was scored at 12 hpi for TIME cells and 24 hpi for U2OS cells using the fixation, staining, and imaging procedures described above. Relative infectivity in KO + EV and KO + PCDH1 cell lines was normalized to the corresponding WT-cell value for each virus.

## Statistics and software

Statistical comparisons, sample size, and exact p-values are reported in the figure legends. ELISA and neutralization data were fit to a four-parameter logistic (4PL) sigmoidal/variable-slope model with log-transformed concentration on the x-axis. All statistical analyses and dose-response curve fitting were performed in GraphPad Prism v11. SnapGene v8.2.2 was used for plasmid design, and phylogenetic analysis was performed in Geneious Prime v2025.2.

## Data availability

No new nucleotide or amino acid sequences, structural data, or custom code were generated in this study. All data supporting these findings are available within the article. Reagents generated in this study are available from the corresponding author upon reasonable request, subject to institutional and biosafety requirements.

## Acknowledgements

Funding for this work was provided by the National Institutes of Health under grants R21AI156482 and R56AI175292 to R.K.J., with additional institutional support from the National Institute of General Medical Sciences through grant P20GM134974 awarded to the Center for Applied Immunology and Pathological Processes (CAIPP). We thank Kartik Chandran for providing recombinant vesicular stomatitis viruses (rVSV), cell lines and antibody-expression plasmids. We also thank Charles M Rice and Francis V Chisari for providing Huh-7.5.1 cells. We are grateful to the Research Core Facility Genomics Core (RRID: SCR_024775), Sushma Bharrhan of the CAIPP Immunophenotyping Core (RRID: SCR_024781), and the CAIPP Bioinformatics and Modeling Core (RRID: SCR_024779) for technical support and access to shared resources. R.K.J. is a co-inventor on a U.S. patent covering methods and assays for treating hantavirus infections.

